# Levels of genetic diversity of SARS-CoV-2 virus: reducing speculations about the genetic variability of the virus in South America

**DOI:** 10.1101/2020.09.14.296491

**Authors:** Pierre Teodósio Felix, Cícero Batista do Nascimento Filho, Robson da Silva Ramos, Antônio João Paulino, Dallynne Bárbara Ramos Venâncio

## Abstract

In this work, we evaluated the levels of genetic diversity in 38 complete Genomes of SARS-CoV-2 from six countries in South America, using specific methodologies for paired F_ST_, AMOVA, mismatch, demographic and spatial expansions, molecular diversity and for the time of evolutionary divergence. The analyses showed non-significant evolutionary divergences within and between the six countries, as well as a significant similarity to the time of genetic evolutionary divergence between all populations. Thus, it seems safe to affirm that we will find similar results for the other Countries of South America, reducing speculation about the existence of rapid and silent mutations that, although there are as we have shown in this work, do not increase, until this moment, the genetic variability of the Virus, a fact that would hinder the work with molecular targets for vaccines and drugs in general.

## 1. Introduction

The new coronavirus, which originated in China, is now expanding into regions marked by poverty, lack of access to water and therefore sanitation and hygiene measures. It takes a geometric scale caused by the complete neglect of the rulers, a very common thing in almost all countries of South America. All these associated factors interfere very much in the epidemiological mechanics of the virus, accentuating the implications of the pandemic. (MILLER *et al*, 2020). Despite this fact, some countries in South America put in place laboratory and patient management protocols used in the SARS outbreak in 2003 and pandemic influenza in 2009, also seeking to establish communication strategies for dissemination of prophylactic measures among neighboring countries, trying to align with what is recommended and recommended by the WHO (RODRIGUEZ-MORALES *et al*., 2020).

From these learned examples, we see the efficient and recent action of the Pan American Health Organization (PAHO/WHO) in the measles outbreak in Latin America, issuing constant epidemiological alerts from January 2019 to January 2020, managing to report 20,430 cases and 19 deaths in 14 countries: Argentina, Bahamas, Brazil, Chile, Colombia, Costa Rica, Cuba, Curacao, Mexico, Peru, Uruguay and Venezuela. However, despite efforts to employ protocols so far efficient in other viruses, in addition to this, the lack of effective drugs against the new coronavirus and vaccines still in the testing phase worldwide, trying to understand the evolutionary aspects of the virus, emerges as another strategy that can help the scientific community discover significant biological aspects of the virus, generating information that can be used, including, in the construction of drugs and vaccines in progress and even in the most varied prophylaxis measures (PAHO, 2020).

Thinking so, the team of the Laboratory of Population Genetics and Computational Evolutionary Biology (LaBECom-UNIVISA) conducted a study of phylogeny and molecular variance analysis to evaluate the possible levels of genetic diversity and polymorphisms existing in a PopSet of the complete genome of SARS-CoV-2 from all over South America, available at the National Center for Biotechnology Information (NCBI), in the Severe acute respiratory syndrome coronavirus 2 data hub (GENBANK, 2020).

## 2. Objective

Evaluate the possible levels of genetic diversity and polymorphism existing in 38 SARS-CoV-2 genomes in South America.

## 3. Methodology

### 3.1. Databank

The 38 complete genome sequences of SARS-CoV-2 from South America (Brazil, Chile, Peru, Colombia, Uruguay and Venezuela with 16, 11, 1, 2, 1, 7 haplotypes, respectively) all with 29,906 pb extension and Phred values ≥ 40 and which now make up our study PopSet, were recovered from GENBANK (https://www.ncbi.nlm.nih.gov/labs/virus/vssi/#/virus?SeqType_s=Nucleotide&VirusLineage_ss=SARS-CoV-2,%20taxid:2697049&Completeness_s=complete&Region_s=South%20America) on August 21, 2020).

### 3.2. Phylogenetic analyses

Nucleotide sequences previously described were used for phylogenetic analyses. The sequences were aligned using the MEGA X program (TAMURA *et al*., 2018) and the gaps were extracted for the construction of phylogenetic trees.

### 3.3. Genetic Structuring Analyses

Paired F_ST_ estimators, Molecular Variance (AMOVA), Genetic Distance, mismatch, demographic and spatial expansion analyses, molecular diversity and evolutionary divergence time were obtained with the Software Arlequin v. 3.5 (EXCOFFIER *et al*., 2005) using 1000 random permutations (NEI and KUMAR, 2000). The F_ST_ and geographic distance matrices were not compared.

## 4. Results

### 4.1. General properties of complete SARS-CoV-2 genome sequences from South America

Of the 38 sequences of the complete genome of SARS-CoV-2 in South America with 29,996 bp extension, the analyses revealed the presence of 75 polymorphic sites and of these, only 71 sites were parsimonious-informative. The graphic representation of these sites could be seen in a logo built with the software WEBLOGO 3. (CROOKS *et al*., 2004), where the size of each nucleotide is proportional to its frequency for certain sites. (Figure 1).

**Figure 1:**
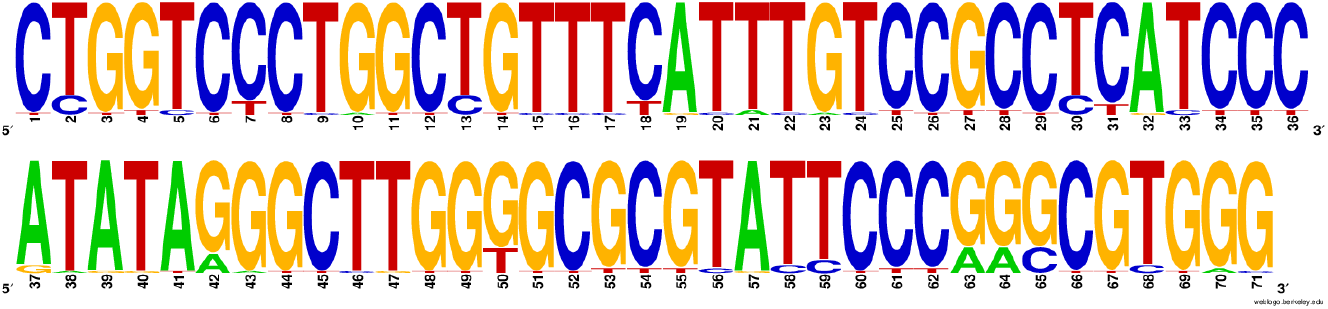
Graphic representation of 71 parsimonium-informative sites of complete Genome Sequences of SARS-CoV-2 from South America.

Using the UPGMA method, based on the 71 parsimony-informative sites, it was also possible to understand that the 38 haplotypes comprised very similar groups genetically and with a non-significant polymorphism pattern (Figure 2).

**Figure 2.**
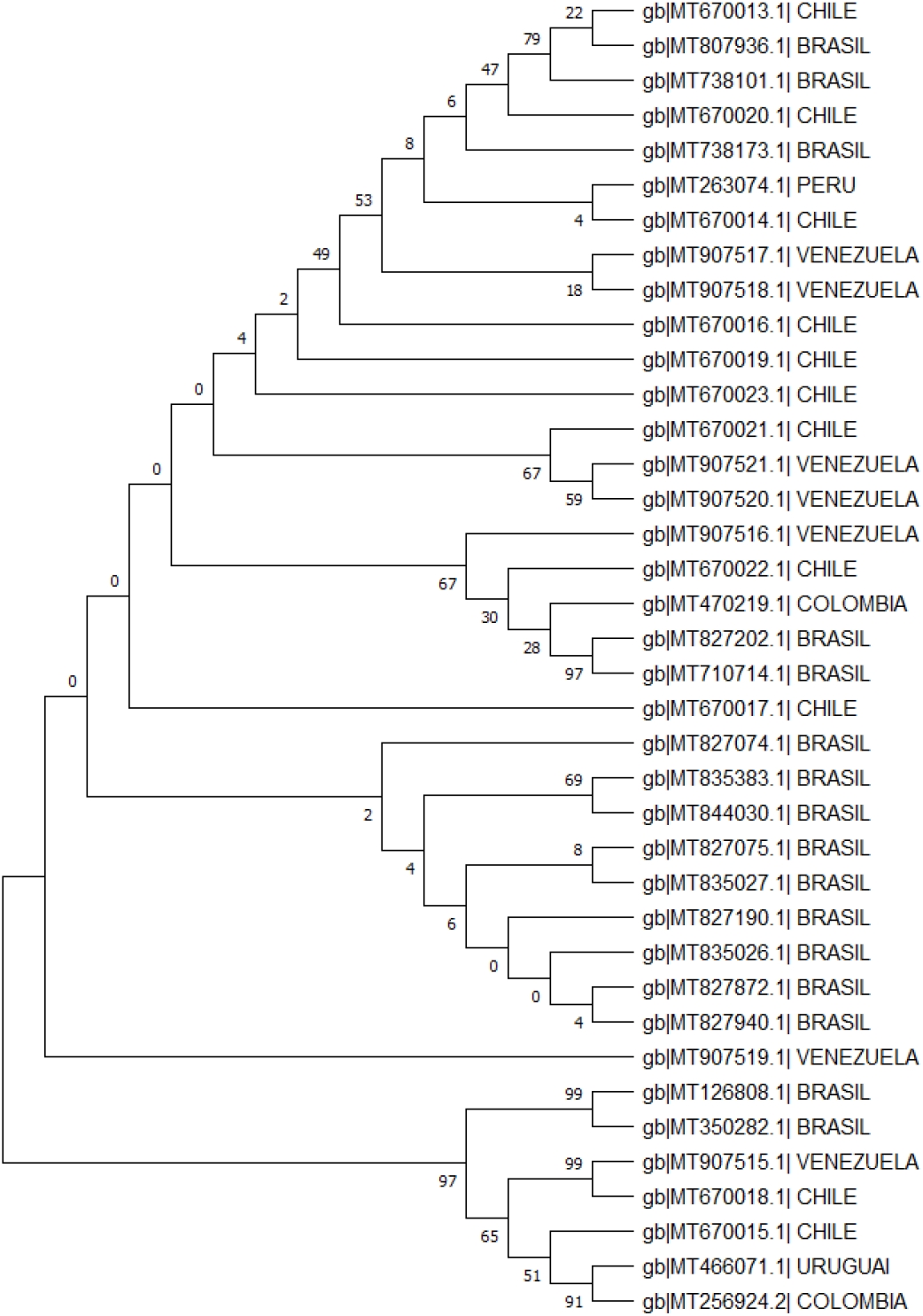
Evolutionary analysis by the maximum likelihood method. The evolutionary history was inferred using the maximum likelihood method and the 3-parameter Tamura model [1]. The tree with the highest probability of logging (−1366.35) is shown. The percentage of trees in which the associated dollar sums group together is shown next to the branches. The initial trees for heuristic research were obtained automatically by applying the Join-Join and BioNJ algorithms to an array of distances in estimated pairs using the Tamura 3 parameter model and then selecting the topology with a higher log probability value. This analysis involved 38 nucleotide sequences. The evolutionary analyses were performed in MEGA X

### 4.2. Molecular Variance Analysis (AMOVA) and Genetic Distance

Genetic distance and molecular variation (AMOVA) analyses were not significant for the groups studied, presenting a variation component of 0.12 between populations and 4.46 within populations. The F_ST_ value (0.03) showed a low fixation index, with non-significant evolutionary divergences within and between groups (Table 1 and Figure 3).

**Table 1.**
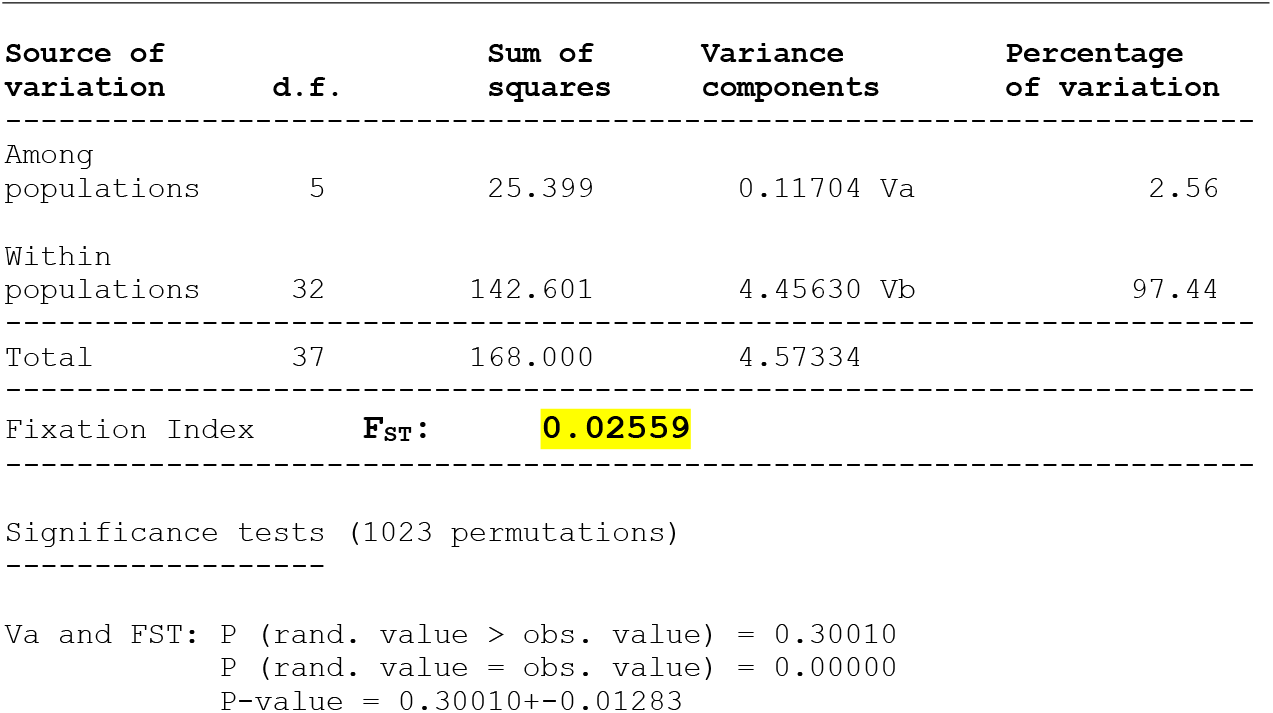
Components of haplotypic variation and paired F_ST_ value for the 38 complete genome sequences of SARS-CoV-2 from South America.

**Figure 3.**
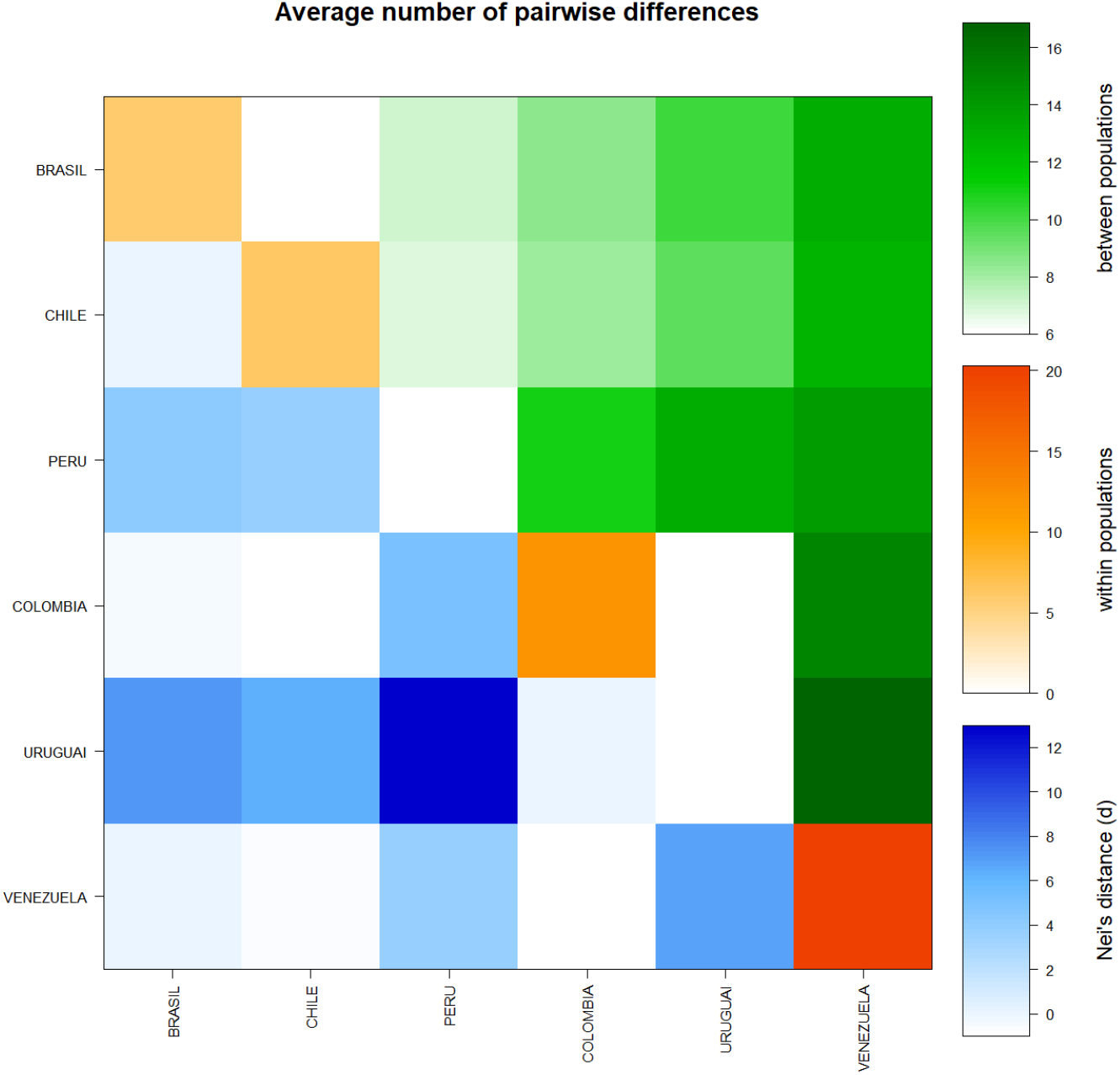
Matrix of paired differences between the populations studied: between the groups; within the groups; and Nei distance for the complete genome sequences of SARS-CoV-2 from six countries in South America.

A significant similarity was also evidenced for the time of genetic evolutionary divergence among all populations; supported by τ variations, mismatch analyses and demographic and spatial expansion analyses. With a representative exception for haplotypes from Brazil and Venezuela (Table 2), (Figures 4 and 5).

**Table 2.** Demographic and spatial expansion simulations based on the τ, θ, and M indices of sequences of the complete SARS-CoV-2 genomes from six South American countries.

| Statistics | BRASIL | CHILE | PERU | COLOMBIA | URUGUAI | VENEZUELA | Mean | s.d. |
| --- | --- | --- | --- | --- | --- | --- | --- | --- |
| **Demographic expansion** |  |  |  |  |  |  |  |  |
| Tau | 8.65821 | 3.41406 | 0.00000 | 0.00000 | 0.00000 | 8.00000 | 3.34538 | 4.08585 |
| Tau qt 2.5% | 1.43937 | 0.00000 | 0.00000 | 0.00000 | 0.00000 | 4.43744 | 0.97947 | 1.78922 |
| Tau qt 5% | 2.76561 | 2.33788 | 0.00000 | 0.00000 | 0.00000 | 5.36523 | 1.74479 | 2.17413 |
| Tau qt 95% | 12.31057 | 17.19734 | 0.00000 | 0.00000 | 0.00000 | 20.92787 | 8.40596 | 9.60534 |
| Tau qt 97.5% | 13.72265 | 18.72268 | 0.00000 | 0.00000 | 0.00000 | 21.95513 | 9.06674 | 10.27271 |
| Theta0 | 0.00000 | 4.28554 | 0.00000 | 0.00000 | 0.00000 | 5.49999 | 1.63092 | 2.55563 |
| Theta0 qt 2.5% | 0.00000 | 0.00000 | 0.00000 | 0.00000 | 0.00000 | 0.00000 | 0.00000 | 0.00000 |
| Theta0 qt 5% | 0.00000 | 0.00000 | 0.00000 | 0.00000 | 0.00000 | 0.00000 | 0.00000 | 0.00000 |
| Theta0 qt 95% | 1.82648 | 3.26617 | 0.00000 | 0.00000 | 0.00000 | 12.13044 | 2.87052 | 4.72678 |
| Theta0 qt 97.5% | 2.83189 | 4.79179 | 0.00000 | 0.00000 | 0.00000 | 16.98018 | 4.10064 | 6.60932 |
| Theta1 | 8.67921 | 13.59864 | 0.00000 | 0.00000 | 0.00000 | 3414.97837 | 572.87604 | 1392.35166 |
| Theta1 qt 2.5% | 3.11981 | 3.91268 | 0.00000 | 0.00000 | 0.00000 | 27.98066 | 5.83553 | 10.98763 |
| Theta1 qt 5% | 4.26270 | 4.56568 | 0.00000 | 0.00000 | 0.00000 | 39.51658 | 8.05749 | 15.56301 |
| Theta1 qt 95% | 33.99159 | 82.50502 | 0.00000 | 0.00000 | 0.00000 | 323.11280 | 73.26823 | 126.61357 |
| Theta1 qt 97.5% | 52.81974 | 162.02985 | 0.00000 | 0.00000 | 0.00000 | 590.61070 | 134.24338 | 232.26574 |
| SSD | 0.04330 | 0.00540 | 0.00000 | 0.00000 | 0.00000 | 0.07165 | 0.02006 | 0.03041 |
| Model (SSD) p-value | 0.30000 | 0.99000 | 0.00000 | 0.00000 | 0.00000 | 0.04000 | 0.22167 | 0.39418 |
| Raggedness index | 0.07035 | 0.00860 | 0.00000 | 0.00000 | 0.00000 | 0.16780 | 0.04112 | 0.06787 |
| Raggedness p-value | 0.38000 | 1.00000 | 0.00000 | 0.00000 | 0.00000 | 0.22000 | 0.26667 | 0.39144 |
| **Spatial expansion** |  |  |  |  |  |  |  |  |
| Tau | 6.18844 | 2.25056 | 0.00000 | 0.00000 | 0.00000 | 8.24067 | 2.77994 | 3.60283 |
| Tau qt 2.5% | 1.28581 | 0.69166 | 0.00000 | 0.00000 | 0.00000 | 3.44275 | 0.90337 | 1.34817 |
| Tau qt 5% | 3.98499 | 1.97354 | 0.00000 | 0.00000 | 0.00000 | 4.37483 | 1.72223 | 2.05513 |
| Tau qt 95% | 10.32285 | 13.44850 | 0.00000 | 0.00000 | 0.00000 | 14.53488 | 6.38437 | 7.12916 |
| Tau qt 97.5% | 10.82249 | 17.56023 | 0.00000 | 0.00000 | 0.00000 | 15.32114 | 7.28398 | 8.26907 |
| Theta | 1.64652 | 4.96606 | 0.00000 | 0.00000 | 0.00000 | 5.15534 | 1.96132 | 2.48474 |
| Theta qt 2.5% | 0.00072 | 0.00072 | 0.00000 | 0.00000 | 0.00000 | 0.00072 | 0.00036 | 0.00040 |
| Theta qt 5% | 0.00072 | 0.00072 | 0.00000 | 0.00000 | 0.00000 | 0.00258 | 0.00067 | 0.00100 |
| Theta qt 95% | 2.75007 | 7.34736 | 0.00000 | 0.00000 | 0.00000 | 16.21983 | 4.38621 | 6.46833 |
| Theta qt 97.5% | 3.02024 | 7.64974 | 0.00000 | 0.00000 | 0.00000 | 20.05757 | 5.12126 | 7.90674 |
| M | 2.30868 | 11.22560 | 0.00000 | 0.00000 | 0.00000 | 8827.30237 | 1473.47278 | 3602.62866 |
| M qt 2.5% | 0.52435 | 0.86722 | 0.00000 | 0.00000 | 0.00000 | 20.59232 | 3.66398 | 8.30087 |
| M qt 5% | 0.82693 | 1.18551 | 0.00000 | 0.00000 | 0.00000 | 33.82173 | 5.97236 | 13.65272 |
| M qt 95% | 12.86742 | 110.88566 | 0.00000 | 0.00000 | 0.00000 | 2097.48882 | 370.20698 | 847.30175 |
| M qt 97.5% | 15.34689 | 191.55404 | 0.00000 | 0.00000 | 0.00000 | 5327.87210 | 922.46217 | 2159.51526 |
| SSD | 0.02288 | 0.00560 | 0.00000 | 0.00000 | 0.00000 | 0.07137 | 0.01664 | 0.02824 |
| Model (SSD) p-value | 0.77000 | 0.98000 | 0.00000 | 0.00000 | 0.00000 | 0.10000 | 0.30833 | 0.44562 |
| Raggedness index | 0.07035 | 0.00860 | 0.00000 | 0.00000 | 0.00000 | 0.16780 | 0.04112 | 0.06787 |
| Raggedness p-value | 0.68000 | 1.00000 | 0.00000 | 0.00000 | 0.00000 | 0.22000 | 0.31667 | 0.42641 |

**Table 3.** Molecular Diversity Indices for the complete Genomes of SARS-CoV-2 from six countries in South America

| Statistics | BRASIL | CHILE | PERU | COLOMBIA | URUGUAI | VENEZUELA | Mean | s.d. |
| --- | --- | --- | --- | --- | --- | --- | --- | --- |
| No. of transitions | **21** | 16 | 0 | 9 | 0 | **28** | 12.333 | 11.396 |
| No. of transversions | 7 | 2 | 0 | 3 | 0 | 14 | 4.333 | 5.391 |
| No. of substitutions | **28** | 18 | 0 | 12 | 0 | **42** | 16.667 | 16.428 |
| No. of indels | 0 | 0 | 0 | 0 | 0 | 0 | 0.000 | 0.000 |
| No. of ts. sites | 21 | 16 | 0 | 9 | 0 | 28 | 12.333 | 11.396 |
| No. of tv. sites | 7 | 2 | 0 | 3 | 0 | 14 | 4.333 | 5.391 |
| No. of subst. sites | 28 | 18 | 0 | 12 | 0 | 42 | 16.667 | 16.428 |
| No. private subst. sites | 20 | 5 | 0 | 4 | 0 | 27 | 9.333 | 11.378 |
| No. of indel sites | 0 | 0 | 0 | 0 | 0 | 0 | 0.000 | 0.000 |
| Pi | 5.942 | 6.236 | 0.000 | 12.000 | 0.000 | 13.143 | 6.22015 | 5.63540 |
| Theta_k | N.A. | N.A. | N.A. | N.A. | N.A. | N.A. | N.A. | N.A. |
| Theta_k_lower | N.A. | N.A. | N.A. | N.A. | N.A. | N.A. | N.A. | N.A. |
| Theta_k_upper | N.A. | N.A. | N.A. | N.A. | N.A. | N.A. | N.A. | N.A. |
| Theta_H | N.A. | N.A. | N.A. | N.A. | N.A. | N.A. | N.A. | N.A. |
| s.d. Theta_H | N.A. | N.A. | N.A. | N.A. | N.A. | N.A. | N.A. | N.A. |
| Theta_S | 8.43824 | 6.14551 | 0.00000 | 12.00000 | 0.00000 | 17.14286 | 7.28777 | 6.75543 |
| s.d. Theta_S | 3.34086 | 2.74879 | 0.00000 | 8.83176 | 0.00000 | 8.00564 | 3.82118 | 3.82620 |
| Theta_pi | 5.94167 | 6.23636 | 0.00000 | 12.00000 | 0.00000 | 13.14286 | 6.22015 | 5.63540 |
| s.d. Theta_pi | 3.35170 | 3.61960 | 0.00000 | 12.49000 | 0.00000 | 7.73070 | 4.53200 | 4.83456 |

**Table 4.** Neutrality Tests for the complete Genomes of SARS-CoV-2 from six countries in South America

|  | Statistics | BRASIL | CHILE | PERU | COLOMBIA | URUGUAI | VENEZUELA | Mean | s.d. |
| --- | --- | --- | --- | --- | --- | --- | --- | --- | --- |
| Ewens-Watterson test |  |  |  |  |  |  |  |  |  |
|  | Sample size | 16 | 11 | 1 | 2 | 1 | 7 | 6.33333 | 6.18601 |
|  | No. of alleles(unchecked) | 16 | 11 | 1 | 2 | 1 | 7 | 6.33333 | 6.18601 |
|  | Observed F value | N.A. | N.A. | N.A. | N.A. | N.A. | N.A. | N.A. | N.A. |
|  | Expected F value | N.A. | N.A. | N.A. | N.A. | N.A. | N.A. | N.A. | N.A. |
|  | Watterson test: Pr(rand F <= obs F) | N.A. | N.A. | N.A. | N.A. | N.A. | N.A. | N.A. | N.A. |
|  | Slatkin's exact test P-value | N.A. | N.A. | N.A. | N.A. | N.A. | N.A. | N.A. | N.A. |
| Chakraborty's test |  |  |  |  |  |  |  |  |  |
|  | Sample size | 16 | 11 | 1 | 2 | 1 | 7 | 6.33333 | 6.18601 |
|  | No. of alleles(unchecked) | 16 | 11 | 1 | 2 | 1 | 7 | 6.33333 | 6.18601 |
|  | Obs. homozygosity | 0.00000 | 0.00000 | 0.00000 | 0.00000 | 0.00000 | 0.00000 | 0.00000 | 0.00000 |
|  | Exp. no. of alleles | 8.13974 | 6.67071 | 0.00000 | 1.92308 | 0.00000 | 5.78902 | 3.75376 | 3.56148 |
|  | P(k or more alleles) | N.A. | N.A. | N.A. | N.A. | N.A. | N.A. | N.A. | N.A. |
| Tajima's D test |  |  |  |  |  |  |  |  |  |
|  | Sample size | 16 | 11 | 1 | 2 | 1 | 7 | 6.33333 | 6.18601 |
|  | S | 28 | 18 | 0 | 12 | 0 | 42 | 16.66667 | 16.42762 |
|  | Pi | 5.94167 | 6.23636 | 0.00000 | 12.00000 | 0.00000 | 13.14286 | 6.22015 | 5.63540 |
|  | Tajima's D | -1.21891 | 0.06649 | 0.00000 | 0.00000 | 0.00000 | -1.34385 | -0.41604 | 0.67194 |
|  | Tajima's D p-value | 0.10600 | 0.57300 | 1.00000 | 1.00000 | 1.00000 | 0.07200 | 0.62517 | 0.44716 |
| Fu's FS test |  |  |  |  |  |  |  |  |  |
|  | No. of alleles(unchecked) | 16 | 11 | 1 | 2 | 1 | 7 | 6.33333 | 6.18601 |
|  | Theta_pi | 5.94167 | 6.23636 | 0.00000 | 12.00000 | 0.00000 | 13.14286 | 6.22015 | 5.63540 |
|  | Exp. no. of alleles | 8.13974 | 6.67071 | 0.00000 | 1.92308 | 0.00000 | 5.78902 | 3.75376 | 3.56148 |
|  | FS | -12.00112 | -6.00361 | 0.00000 | 2.48491 | 0.00000 | -1.09653 | -2.76939 | 5.31846 |
|  | FS p-value | 0.00000 | 0.00200 | N.A. | 0.56600 | N.A. | 0.18400 | N.A. | N.A. |

**Figure 4.**
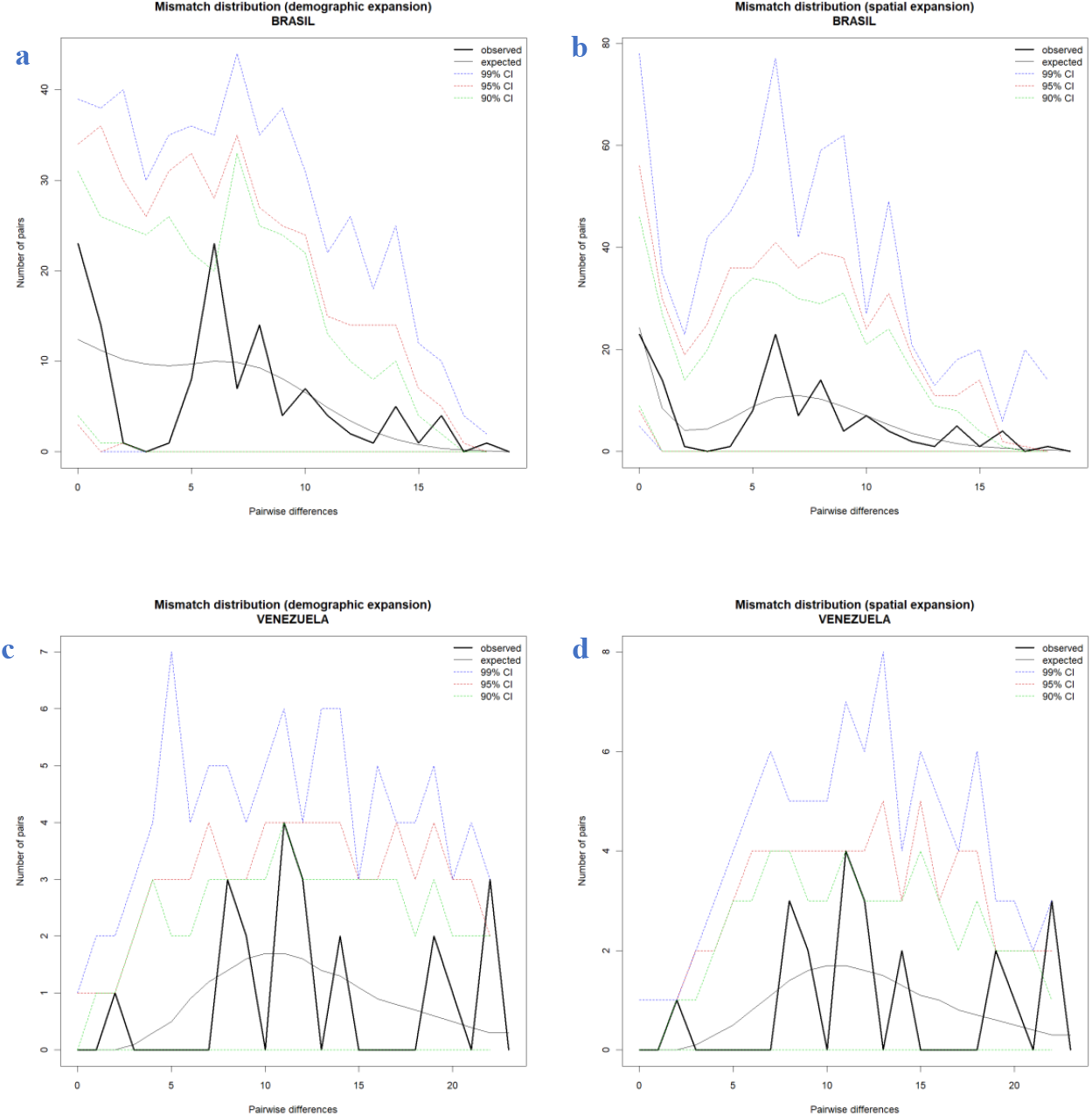
Comparison between the Demographic and Spatial Expansion of sequences of the complete genomes of SARS-CoV-2 from six countries in South America. (**a** and **b**) Graphs of demographic expansion and spatial expansion of haplotypes from Brazil, respectively; (**c** and **d**) Graphs of demographic expansion and spatial expansion of haplotypes from Venezuela, respectively. *Graphs Generated by the statistical package in R language using the output data of the Software Arlequin version 3.5.1.2

**Figure 5.**
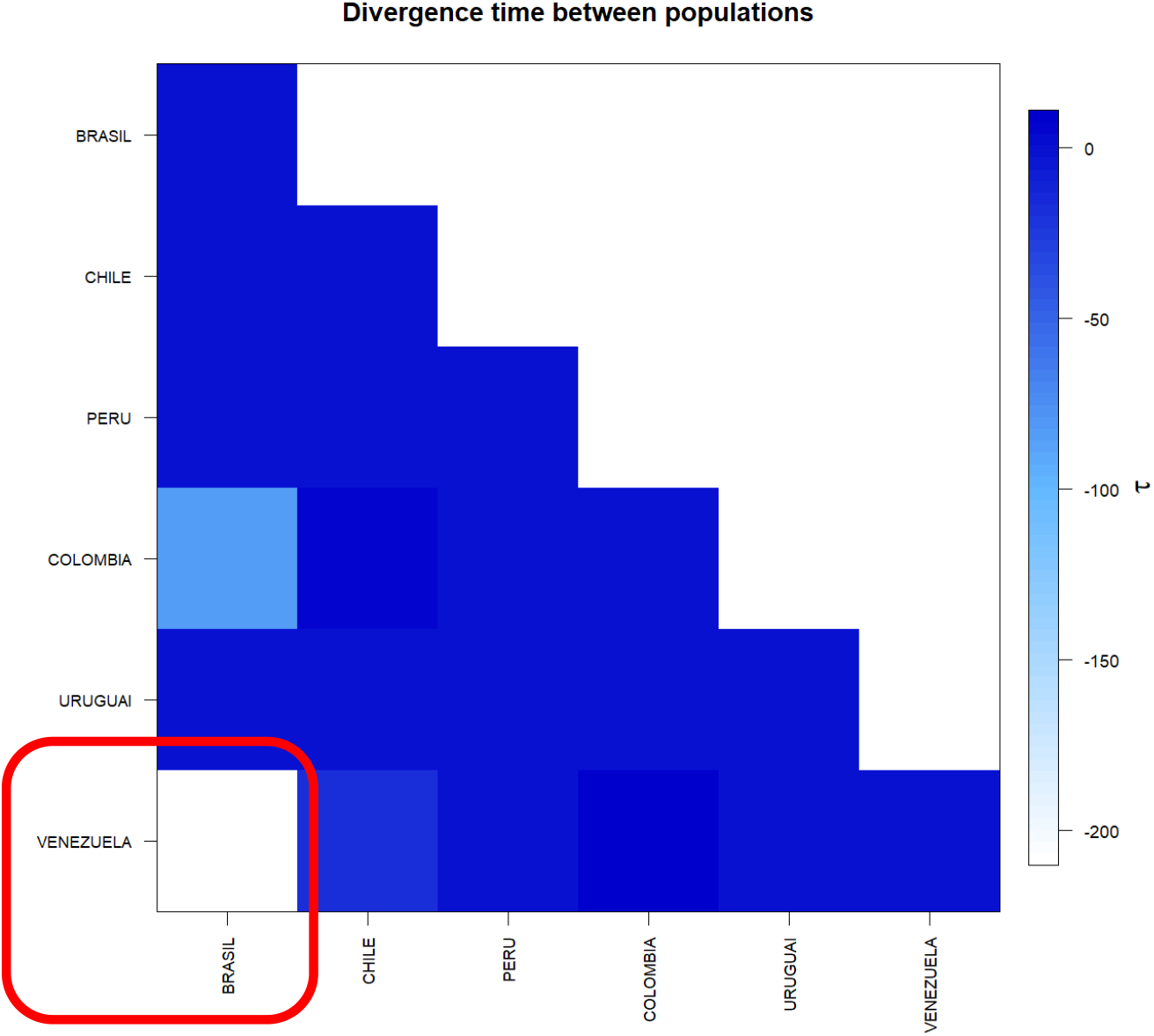
Matrix of divergence time between the complete genomes of SARS-CoV-2 from six countries in South America. In evidence the high value τ present between the sequences of Brazil and Venezuela. * Generated by the statistical package in R language using the output data of the Software Arlequin version 3.5.1.2.

The molecular diversity analyses estimated per θ reflected a significant level of mutations among all haplotypes (transitions and transversions). Indel mutations (insertions or additions) were not found in any of the six groups studied. The D tests of Tajima and Fs de Fu showed disagreements between the estimates of general θ and π, but with negative and highly significant values, indicating, once again, an absence of population expansions. The irregularity index (R= Raggedness) with parametric bootstrap, simulated new θ values for before and after a supposed demographic expansion and in this case assumed a value equal to zero for all groups (Table 6); (Figure 6).

**Figure 6.**
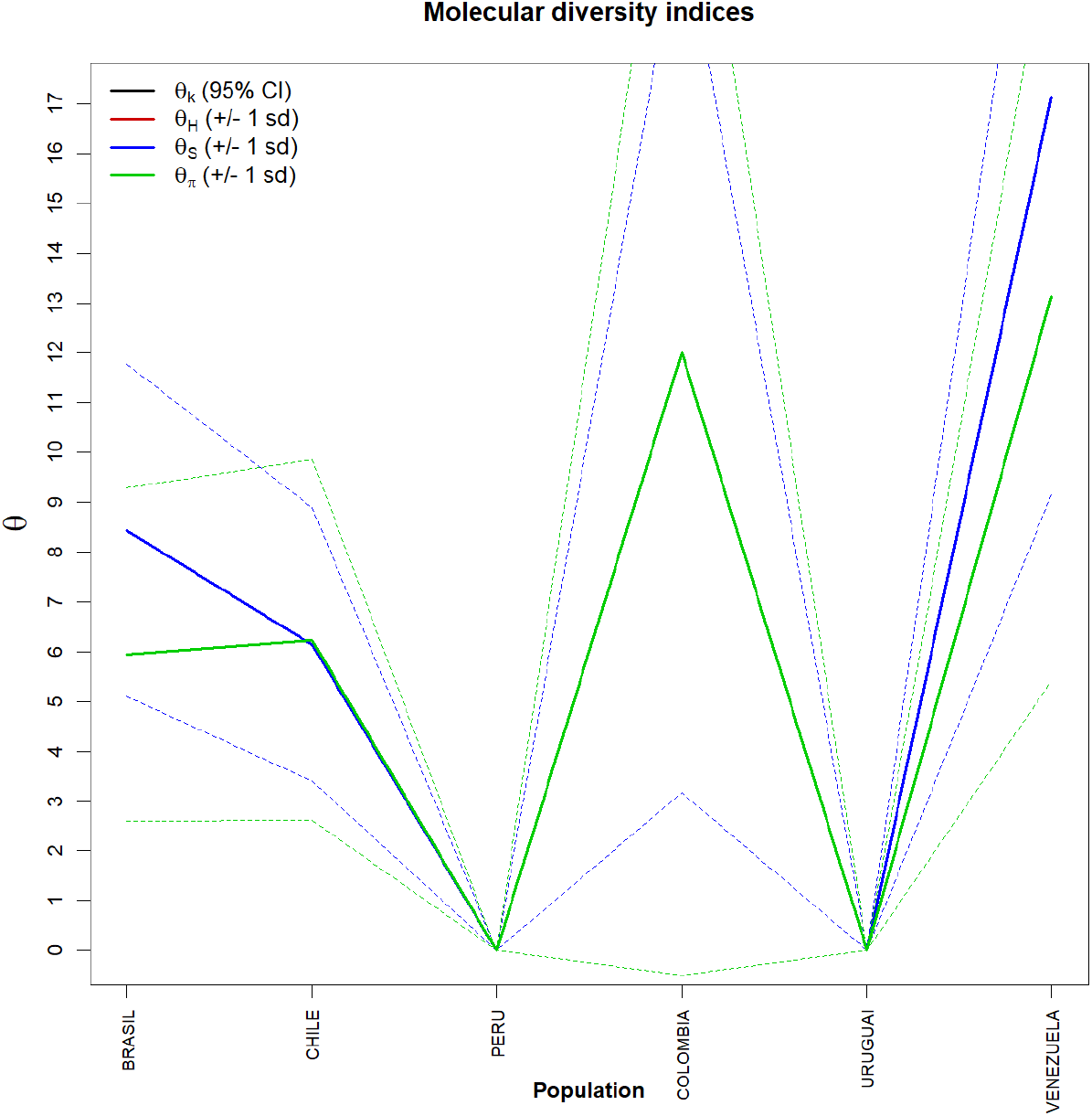
Graph of molecular diversity indices for the complete genomes of SARS-CoV-2 from six countries in South America. In the graph the values of θ: (θk) Relationship between the expected number of alllos (k) and the sample size; (θH) Expected homozygosity in a balanced relationship between drift and mutation; (θS) Relationship between the number of segregating sites (S), sample size (n) and non-recombinant sites; (θπ) Relationship between the average number of paired differences (π) and θ. * Generated by the statistical package in R language using the output data of the Arlequin software version 3.5.1.2.

## 5. Discussion

As the use of phylogenetic analysis and population structure methodologies had not yet been used in this PopSet, in this study it was possible to detect the existence of 6 distinct groups for the complete genome sequences of SARS-CoV-2 from South America, but with minimal variations among all of them. The groups described here presented minimum structuring patterns and were effectively slightly higher for the populations of Brazil and Venezuela. These data suggest that the relative degree of structuring present in these two countries may be related to gene flow. These structuring levels were also supported by simple phylogenetic pairing methodologies such as UPGMA, which in this case, with a discontinuous pattern of genetic divergence between the groups (supports the idea of possible sub-geographical isolations resulting from past fragmentation events), was observed a not so numerous amount of branches in the tree generated and with few mutational steps.

These few mutations have possibly not yet been fixed by drift by the lack of the founding effect, which accompanies the behavior of dispersion and/or loss of intermediate haplotypes throughout the generations. The values found for genetic distance support the presence of this continuous pattern of low divergence between the groups studied, since they considered important the minimum differences between the groups, when the haplotypes between them were exchanged, as well as the inference of values greater than or equal to that observed in the proportion of these permutations, including the p-value of the test.

The discrimination of the 38 genetic entities in their localities was also perceived by their small inter-haplotypic variations, hierarchised in all covariance components: by their intra- and inter-individual differences or by their intra- and intergroup differences, generating a dendogram that supports the idea that the significant differences found in countries such as Brazil and Venezuela, for example, were shared more in their form than in their number, since the result of estimates of the average evolutionary divergence found within these and other countries, even if they exist, were very low.

Based on the high level of haplotypic sharing, tests that measure the relationship between genetic distance and geographic distance, such as the Mantel test, were dispensed in this Estimators θ, even though they are extremely sensitive to any form of molecular variation (FU, 1997), supported the uniformity between the results found by all the methodologies employed, and can be interpreted as a phylogenetic confirmation that there is a consensus in the conservation of the SARS-CoV-2 genome in the Countries of America of America of South objects of this study, being therefore safe to affirm that the small number of existing polymorphisms should be reflected even in all their protein products. This consideration provides the safety that, although there are differences in the haplotypes studied, these differences are minimal in geographically distinct regions and thus it seems safe to extrapolate the levels of polymorphism and molecular diversity found in the samples of this study to other genomes of other South American countries, reducing speculation about the existence of rapid and silent mutations that, although they exist as we have shown in this work, they can significantly increase the genetic variability of the Virus, making it difficult to work with molecular targets for vaccines and drugs in general.

## STATEMENT

Me and the other authors of the manuscript “Molecular variance analysis (AMOVA) and levels of genetic diversity of complete genome of SARS-CoV-2 virus from of six South American Countries” **declare that there are no competing interests**. We are all from the same laboratory and the work was carried out together.

Pierre Teodosio Felix, Head of research, Laboratory of Population Genetics and Computational Evolutionary Biology - LaBECom, UNIVISA, Vitória de Santo Antão, Pernambuco

